# Incomplete Dominance of *ASIP* Alleles in Hungarian Puli Dogs is Associated with *MC1R* Mutation

**DOI:** 10.64898/2026.03.17.712399

**Authors:** Stepan N. Belyakin, Daniil A. Maksimov, Maria A. Pobedintseva, Petr P. Laktionov, Nadezhda V. Mikhnevich, Fedor A. Sipin, Mariya I. Krylova

## Abstract

Alleles of *ASIP* gene (Agouti locus) in dogs determine a wide spectrum of coat colors, from red to black. Gain-of-function *Ay* allele is the most dominant in the range of known *ASIP* mutations: when all other genes affecting coat pigmentation are intact, presence of *Ay* allele results in red coat color. Loss-of-function *a* allele is the most recessive allele of this gene. When homozygous, it gives black coat color. Usually, dogs with *Ay/a* genotype have red coat, because a single copy of *Ay* allele is sufficient to fully compensate for the non-functional allele *a*, implying the complete dominance in this pair of alleles. However exceptions are known. In the Hungarian Puli breed there is a specific coat pigmentation type called fakó. We investigated the genetic composition of fakó dogs and found evidence that the dominance of the *Ay* allele over the *a* allele may be incomplete in these dogs. Analysis of the *MC1R* gene that interacts with *ASIP* in the hair pigmentation genetic cascade allowed us to find the variants that may be responsible for the incomplete dominance of *Ay* allele over *a* allele in Hungarian Puli dogs.

## Introduction

In dogs, coat pigment production in melanocytes is controlled by several genes, among which *ASIP* and *MC1R* play a central role [1,2]. The *MC1R* gene encodes the melanocortin 1 receptor, a transmembrane protein located on the surface of melanocytes. In the presence of MSH (melanocyte-stimulating hormone), *MC1R* activates the synthesis of eumelanin, the dark pigment of hair. *ASIP* (agouti signaling protein) antagonizes MSH and switches melanocytes to produce pheomelanin, a lighter yellowish-red pigment typical of red hair [1,3,4].

Several mutations are known in both the *ASIP* and *MC1R* genes, some of which have been associated with specific phenotypes [5-13]. A recent study elucidated the molecular basis of the known *ASIP* alleles. It was shown that the *Ay* (dominant red), *Ays* (shaded red), *aw* (wild type), *asa* (saddle), and *at* (tan points) alleles are characterized by distinct promoter variants that cause differences in *ASIP* expression and mediate the corresponding phenotypes [12]. In contrast, the *a* (recessive black) allele is regarded as a loss-of-function mutation in the protein-coding region of the *ASIP* gene [13]. Accordingly, the dominance hierarchy of *ASIP* alleles approximately follows the order: *Ay*>*Ays*>*aw*>*asa*=*at*>*a* [12,14].

There are known cases of incomplete dominance within the hierarchy of *ASIP* alleles. One well-known example concerns the interaction between the *asa* and *at* alleles, where *asa*/*at* compound heterozygotes exhibit an intermediate phenotype between the saddle pattern and tan points [15]. More recently, it was shown that the rare sesame coat colors in Shiba Inu are caused by incomplete dominance of the *Ays* or *aw* alleles over *at* (genotypes *Ays*/*at* or *aw*/*at*) [16].

The *MC1R* gene also has several alleles associated with different effects on coat pigmentation [5-11]. Some known mutations in *MC1R* have not yet been attributed to any specific phenotype, leaving room for further studies [17]. In this context, a particularly interesting study associated a long-known mutation in the *MC1R* gene with an unusual partial inhibition of eumelanin pigmentation. This ancient hypomorphic allele, termed *eA*, causes the domino coat pattern in northern dog breeds and produces similar effects in many others [9,18]. This example suggests that other known mutations in the *MC1R* gene may also influence hair pigmentation in dogs in ways that are not yet fully understood.

Here, we investigated an unusual pigmentation pattern in the Hungarian Puli breed, the inheritance of which has long puzzled breeders. This pattern, referred to as fakó, is characterized by a darker eumelanin overlay on a fawn background. The pattern is illustrated in the Figure 1 (first column, “shaded”). The second column (“non-shaded”) shows red (fawn) dogs with or without a melanistic mask.

**Figure 1.**
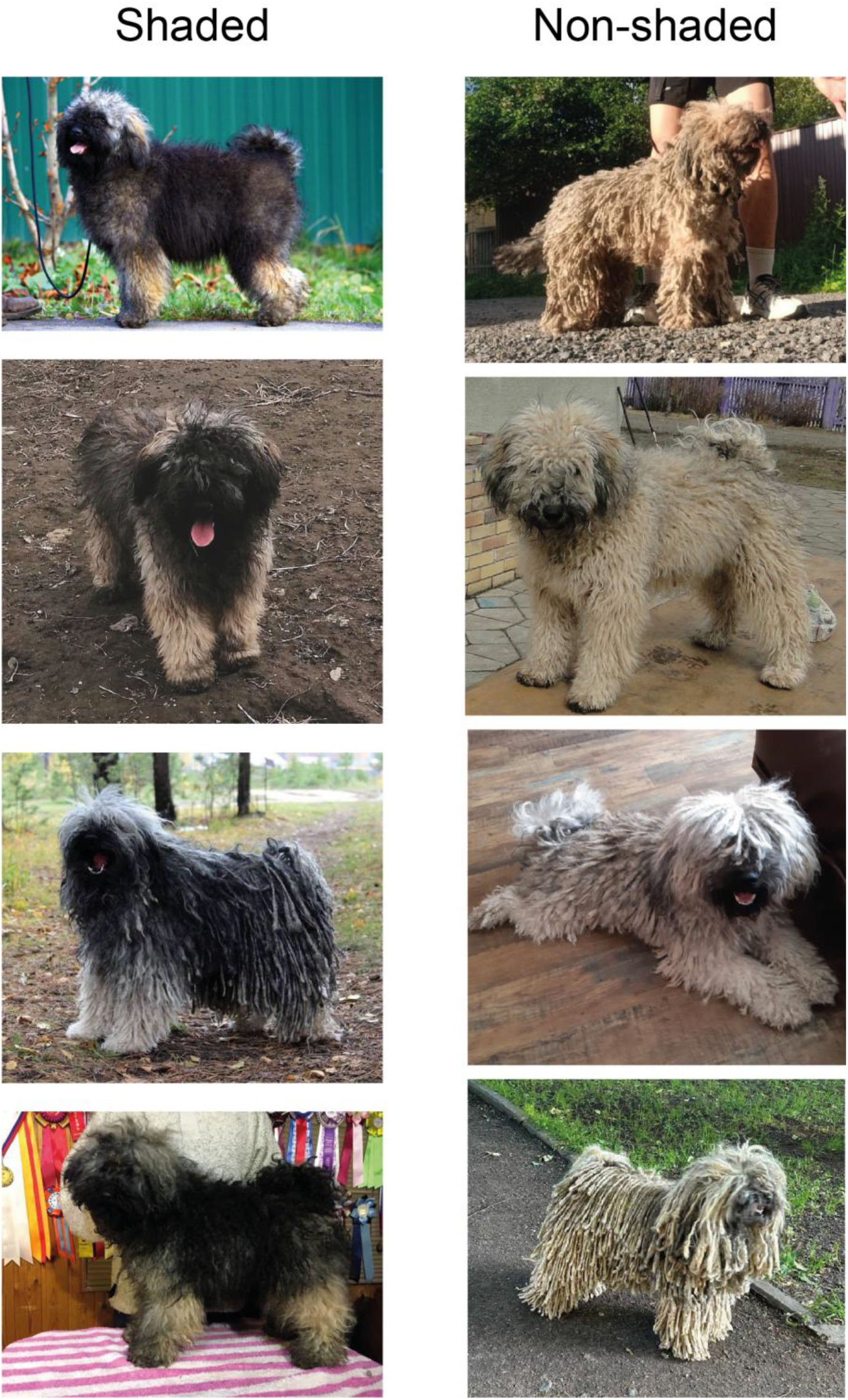
Shaded and non-shaded Puli dogs. All dogs presented represent *Ay*/*a* genotype in *ASIP* gene (details in the text).

Existing data indicate that three *ASIP* alleles are present in the Puli breed: *Ay, at*, and *a* [19]. The *MC1R* gene in this breed is represented by the *Em* allele (melanistic mask), the wild-type *E* allele, and the recessive red *e* allele [19]. However, these alleles are not sufficient to explain the shaded phenotype observed in this breed. Therefore, we genotyped the *ASIP* and *MC1R* genes in shaded and non-shaded Puli dogs to investigate the genetic basis of the fakó pigmentation pattern.

## Results

For this study, we collected samples from shaded and non-shaded fawn Puli dogs provided by breeders and owners in Russia. The dogs were phenotyped based on photographs submitted together with buccal swabs. The final set consisted of 12 phenotypically shaded dogs and 11 non-shaded dogs.

All dogs were genotyped for the known alleles of the *ASIP* gene (see Materials and Methods). Remarkably, all 12 dogs with the shaded fawn phenotype carried the *Ay*/*a* genotype (Table 1). Importantly, our genotyping approach distinguished between the *Ay* and *Ays* alleles [12,16], and all dogs in this group carried the most dominant *Ay* allele, indicating that the mechanism underlying dark shading in these dogs differs from that observed in sesame Shiba Inu [16]. The involvement of the dominant black Kb allele of the CBD103 gene [20], which could potentially contribute to black pigmentation, was ruled out, as none of the shaded dogs carried this allele (data not shown).

**Table 1.**
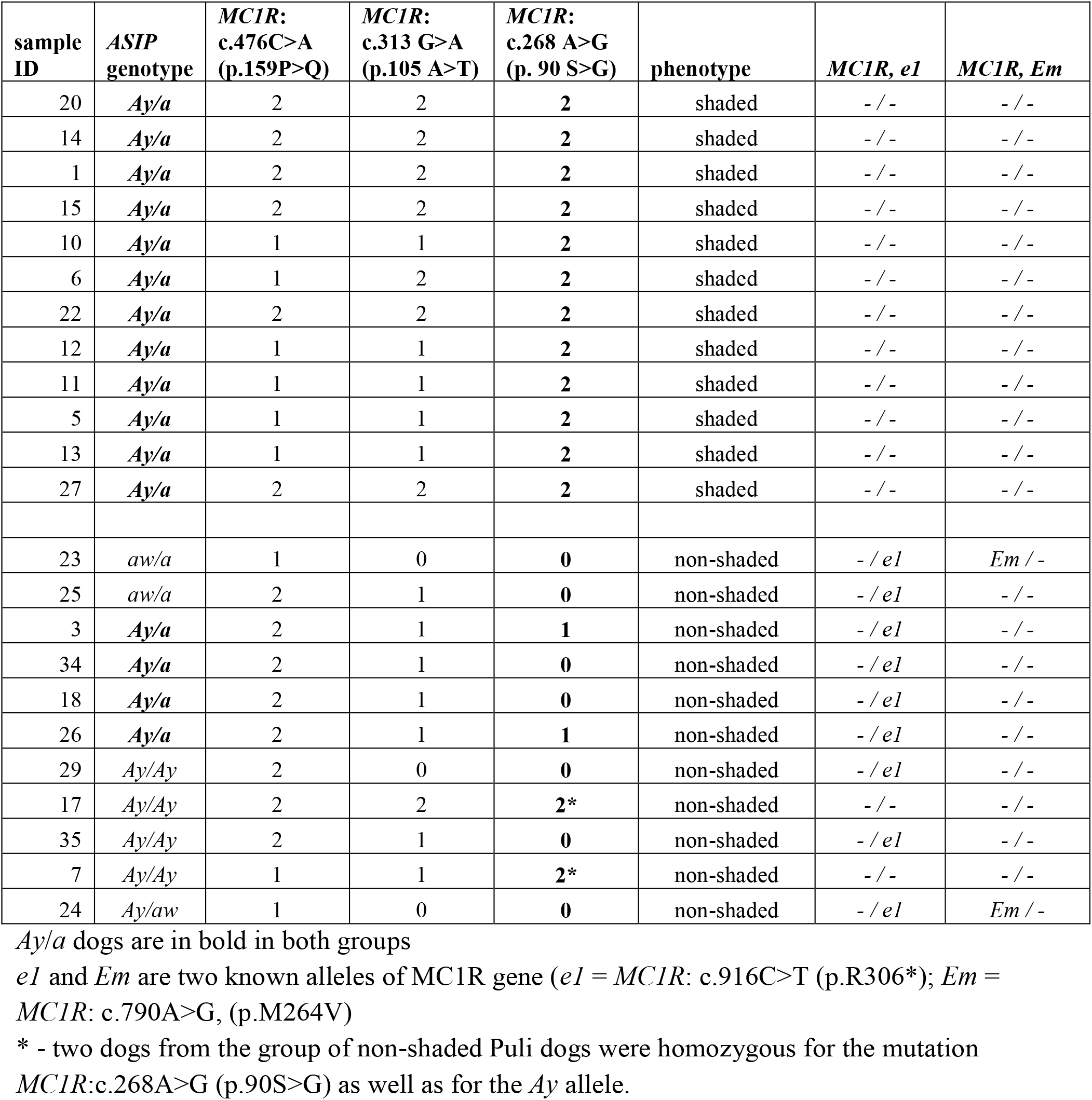
*ASIP* and *MC1R* genotypes of the shaded and non-shaded Puli dogs.

| sample ID | *ASIP* genotype | *MC1R*: c.476C>A (p.159P>Q) | *MC1R*: c.313 G>A (p.105 A>T) | *MC1R*: c.268 A>G (p. 90 S>G) | phenotype | *MC1R, e1* | *MC1R, Em* |
| --- | --- | --- | --- | --- | --- | --- | --- |
| 20 | **Ay/a** | 2 | 2 | **2** | shaded | - / - | - / - |
| 14 | **Ay/a** | 2 | 2 | **2** | shaded | - / - | - / - |
| 1 | **Ay/a** | 2 | 2 | **2** | shaded | - / - | - / - |
| 15 | **Ay/a** | 2 | 2 | **2** | shaded | - / - | - / - |
| 10 | **Ay/a** | 1 | 1 | **2** | shaded | - / - | - / - |
| 6 | **Ay/a** | 1 | 2 | **2** | shaded | - / - | - / - |
| 22 | **Ay/a** | 2 | 2 | **2** | shaded | - / - | - / - |
| 12 | **Ay/a** | 1 | 1 | **2** | shaded | - / - | - / - |
| 11 | **Ay/a** | 1 | 1 | **2** | shaded | - / - | - / - |
| 5 | **Ay/a** | 1 | 1 | **2** | shaded | - / - | - / - |
| 13 | **Ay/a** | 1 | 1 | **2** | shaded | - / - | - / - |
| 27 | **Ay/a** | 2 | 2 | **2** | shaded | - / - | - / - |
| 23 | aw/a | 1 | 0 | **0** | non-shaded | - / e1 | Em / - |
| 25 | aw/a | 2 | 1 | **0** | non-shaded | - / e1 | - / - |
| 3 | **Ay/a** | 2 | 1 | **1** | non-shaded | - / e1 | - / - |
| 34 | **Ay/a** | 2 | 1 | **0** | non-shaded | - / e1 | - / - |
| 18 | **Ay/a** | 2 | 1 | **0** | non-shaded | - / e1 | - / - |
| 26 | **Ay/a** | 2 | 1 | **1** | non-shaded | - / e1 | - / - |
| 29 | Ay/Ay | 2 | 0 | **0** | non-shaded | - / e1 | - / - |
| 17 | Ay/Ay | 2 | 2 | **2\*** | non-shaded | - / - | - / - |
| 35 | Ay/Ay | 2 | 1 | **0** | non-shaded | - / e1 | - / - |
| 7 | Ay/Ay | 1 | 1 | **2\*** | non-shaded | - / - | - / - |
| 24 | Ay/aw | 1 | 0 | **0** | non-shaded | - / e1 | Em / - |

*Ay/a* dogs are in bold in both groups
*e1* and *Em* are two known alleles of *MC1R* gene (*e1* = *MC1R*: c.916C>T (p.R306\*); *Em* = *MC1R*: c.790A>G, (p.M264V)
\* - two dogs from the group of non-shaded Puli dogs were homozygous for the mutation *MC1R*:c.268A>G (p.90S>G) as well as for the *Ay* allele.

Non-shaded dogs possessed the following combinations of *ASIP* alleles: *Ay*/*Ay* (4 dogs), *Ay*/*aw* (1 dog), *aw*/*a* (2 dogs), and *Ay*/*a* (4 dogs) (Table 1). The phenotypes of dogs with the *Ay*/*Ay, Ay*/*aw*, and *aw*/*a* genotypes accurately corresponded to the known dominance hierarchy of *ASIP* alleles (*Ay*>*aw*>*a*), with *Ay* being the most dominant allele and *a* the most recessive. Accordingly, these non-shaded dogs exhibited either red (fawn) or wolf-sable coat colors, as exemplified in the Figure 1). However, four dogs in this group carried the *Ay*/*a* genotype—the same genotype observed in shaded dogs—but did not display a dark overlay (Table 1). As in the shaded group, none of the dogs in this group carried the dominant black allele of the CBD103 gene (data not shown).

The *Ay* allele is characterized by a combination of the two strongest promoters known in the *ASIP* gene [12]. This allele produces a high level of *ASIP* protein, leading to sustained occupation of the *MC1R* receptor on the surface of melanocytes and resulting in uniform red coat pigmentation. In contrast, the *a* allele does not produce functional *ASIP* protein capable of binding the *MC1R* receptor [13]. In our study the non-shaded *Ay*/*a* dogs illustrate the complete dominance of *Ay* over the *a* allele. In contrast, in shaded Puli dogs with the *Ay*/*a* genotype, a single copy of the *Ay* allele appears to be insufficient to fully mask the presence of the loss-of-function *a* allele, suggesting that the dominance of *Ay* over *a* is altered in these animals.

*MC1R* interacts with the *ASIP* protein, and alleles of the *MC1R* gene are known to modify phenotypes produced by *ASIP* alleles [5,7,9,10]. Therefore, we analyzed the *MC1R* gene in both groups. We performed Sanger sequencing of the entire coding sequence (CDS) to identify alleles with known phenotypic effects as well as genetic variants that had not previously been associated with coat color variation.

In the group of shaded dogs, we observed three variants that had not previously been associated with any phenotype: c.476C>A (p.159P>Q), c.313G>A (p.105A>T), and c.268A>G (p.90S>G) [17]. Among these variants, only c.268A>G (p.90S>G) was homozygous in all shaded dogs. Variant c.313G>A (p.105A>T) was heterozygous in three dogs. Variant c.476C>A (p.159P>Q) was heterozygous in three dogs and absent in two dogs. Interestingly, in five dogs all three variants were present in the homozygous state, indicating that they may be linked in Puli dogs.

Among non-shaded dogs with the *ASIP* genotype *Ay*/a, none were homozygous for the c.268A>G (p.90S>G) variant. All dogs in this group were heterozygous for c.313G>A (p.105A>T) and homozygous for c.476C>A (p.159P>Q). Thus, among these three variants, c.268A>G (p.90S>G) was homozygous in all shaded dogs and was not present in the homozygous state in non-shaded dogs. The other two variants were detected in different combinations in both groups. In the non-shaded group, we also detected the well-known *MC1R* allele e1: all four dogs were carriers of this allele.

We checked if the variant c.268A>G (p.90S>G) is present in other breeds using available genomic data. Out of 166 analyzed breeds this allele was found in 95 including Newfoundland, Siberian Husky, Tibetan Mastiff, Sheltie and others (Supplementary Table 1). Unfortunately, due to the absence of the phenotypic data we were not able to match the presence of this allele to the coat colors.

Taken together, our results suggest that the shaded phenotype in the Puli breed may be explained by incomplete dominance of the *Ay* allele over the *a* allele of the *ASIP* gene. This incomplete dominance could be caused by the presence of the c.268A>G (p.90S>G) variant, which may reduce the sensitivity of *MC1R* to the *ASIP* ligand and thereby weaken its ability to switch the type of pigment produced in melanocytes.

## Discussion

Incomplete dominance of one allele over another can be caused by different factors. One of the most frequently observed mechanisms is haploinsufficiency, in which a single copy of a dominant allele does not produce sufficient product to fully reconstitute the phenotype. In such cases, the amount of product from the dominant allele is lower than required for full development of the corresponding trait. The sesame pattern in Shiba Inu illustrates how the *Ays* allele of the *ASIP* gene cannot completely mask the presence of the *at* allele [16]. Similarly, the *asa* allele is not fully dominant over *at*, and *asa*/*at* compound genotypes were shown to exhibit an intermediate phenotype between the saddle pattern and tan points [15].

Another possible mechanism is epistasis, in which the manifestation of a dominant allele is affected by other genes. Our results suggest that in Puli dogs, the dominance of the *Ay* allele of the *ASIP* gene may be compromised by a previously uncharacterized variant of the *MC1R* gene.

*Ay* is the strongest allele of the *ASIP* gene in dogs. Usually, a single copy of this allele is sufficient to completely mask the presence of other *ASIP* alleles, producing a solid red coat color due to a high concentration of *ASIP* protein that remains consistently above the threshold required to switch *MC1R* signaling [13,14]. Thus, the *Ay* allele is normally fully dominant over all other *ASIP* alleles.

In shaded Puli dogs, the *Ay* allele appears to be unable to switch *MC1R* signaling as efficiently as it normally does. This phenomenon may have several possible explanations. For example, it may imply the existence of an additional *ASIP* allele that carries the defining markers of the *Ay* allele but has reduced functional strength. Alternatively, a mutation affecting gene(s) involved in the regulation of *ASIP* gene activity may be present. Also, one could hypothesize the presence of a mutation in a completely different, previously uncharacterized gene involved in coat pigmentation. Other possible scenarios could be proposed.

In this study, we explored one such possibility. A mutation in the *MC1R* gene could reduce the ability of the *ASIP* protein to modulate pigment synthesis in melanocytes. Homozygosity for the c.268 A>G (p.90S>G) mutation perfectly correlated with the shaded phenotype in Puli dogs, strongly suggesting that this mutation contributes to the incomplete dominance of the *Ay* allele over the *a* allele. Furthermore, the c.268 A>G (p.90S>G) mutation is located within the same transmembrane domain as the *Eg* [5] and *Eh* [7] alleles, which cause similar shaded phenotypes in other dog breeds. It is therefore plausible that the c.268 A>G (p.90S>G) variant affects *MC1R* function in a comparable manner.

Our analysis showed that this allele is present in many different breeds. It implies that this mutation in *MC1R* gene may also affect the coat pigmentation outside the Hungarian Puli breed. It is also possible that it could similarly modify the coat colors produced by different combinations of other *ASIP* alleles. This would be an interesting extension of this study.

It should be noted that possible alternative explanations were not addressed in this study. Unfortunately, due to the traditional breeding scheme in the Puli breed—where fawn and shaded dogs are usually mated with black-coated partners (which may be either homozygous for the *a* allele or carriers of the dominant black allele of the CBD103 gene)—familial analysis of the inheritance of the shaded pattern is strongly impeded. Such analysis could otherwise be useful for testing at least some of the alternative hypotheses.

Nevertheless, despite the limited number of available samples, our study demonstrates an association between the shaded phenotype in Puli dogs and a specific combination of mutations in the *ASIP* and *MC1R* genes, and provides a plausible explanation for this coat color pattern through an epistatic interaction between the c.268 A>G (p.Ser90Gly) mutation in the *MC1R* gene and the *Ay* allele of the *ASIP* gene.

## Materials and methods

### Samples

All the samples for this study accompanied with photographs of each dog were sent to VetGenomics lab (Novosibirsk, Russia) by the owners with the explicit agreement for their use in this study and anonymous publication of the photographs. Genomic DNAs were isolated from the buccal swabs and used in PCR (BioMaster HS-Taq PCR-Color (2×) mastermix, Biolabmix, Russian Federation). Genotypes were determined by assessing the lengths of PCR products in 1% agarose gel in 1x Tris-Acetate-EDTA buffer.

### ASIP genotyping

The primer sequences and protocol for allele discrimination in locus A were taken from [12]. Allele nomenclature used in this study follows the traditional naming of the ASIP alleles that is familiar to dog breeders and owners [16]

### MC1R analysis

The following primers were used for amplification and sequencing of the MC1R CDS:

MC1R-F: 5’-CCTCACCAGGAACATAGCAC-3’

MC1R-R: 5’-CTGAGCAAGACACCTGAGAG-3’

MC1R-seq1: 5’-GCATTTTCTTTCTCTGCTGGG-3’

MC1R-seq2: 5’-CATTGCCAAGAACCGCAAC-3’

Genomic DNAs were isolated from the buccal swabs using the conventional in-house spin-column based extraction method and used in PCR (BioMaster HS-Taq PCR-Color (2×) mastermix, Biolabmix, Russian Federation) according to the manufacturer’s recommendations. Resulting products were purified using the precipitation with polyethylene glycol [21]. Sequencing reactions were performed with BrightDye terminators (Molecular Cloning Laboratories, BDT3-100) and analyzed on Applied Biosystems 3130 Genetic Analyzer.

### Bioinformatic analysis

Sequencing data from several public bioprojects available in the NCBI BioProject database were used to assess the presence of the *MC1R*:c.268 A>G (p.Ser90Gly) mutation: PRJNA648123, PRJNA961733, PRJNA448733, and PRJNA288568 [22-25]. The full dataset included 1053 samples representing 164 breeds of domestic dogs (*Canis lupus familiaris*), as well as samples from wolves (*Canis lupus*) and the coyote (*Canis latrans*).

Preprocessing and quality control of raw reads were performed using fastp v0.22.0 [26]. Filtered reads were mapped to the dog reference genome (CanFam3.1) using BWA-MEM2 v2.2.1 [27,28]. Variant calling and subsequent processing were carried out using the Genome Analysis Toolkit (GATK) v4.6.2.0 according to the developers’ best-practice recommendations [29].

## Conflict of interests

SNB, DAM, MAP, PPL, and NVM are employed at the VetGenomics Company (Novosibirsk, Russia), which provides commercial genetic tests for domestic animals.

## Acknowledgements

Authors are grateful to all Hungarian Puli breeders and owners who agreed to participate in this study.

